# Human herpesvirus 1 associated with epizootics in Belo Horizonte, Minas Gerais, Brazil

**DOI:** 10.1101/2025.03.24.645062

**Authors:** Gabriela F. Garcia-Oliveira, Mikaelly Frasson Biccas, Daniel Jacob, Marcelle Alves de Oliveira, Ana Maria de Oliveira Paschoal, Pedro A. Alves, Cecília Barreto, Daniel Ambrósio da Rocha Vilela, Érika Procópio Tostes Teixeira, Thiago Lima Stehling, Thais Melo Mendes, Marlise Costa Silva, Munique Guimarães Almeida, Ivan Vieira Sonoda, Érica Munhoz de Mello, Francisco Elias Nogueira da Gama, Kathryn A. Hanley, Nikos Vasilakis, Betânia Paiva Drumond

## Abstract

Human activity in sylvatic environments and resulting contact with wildlife, such as non-human primates (NHP), can lead to pathogen spillover or spillback. Both NHPs and humans host a variety of herpesviruses. While these viruses typically cause asymptomatic infections in their natural hosts, they can lead to severe disease or even death when they move into novel hosts. In early 2024, deaths of *Callithrix penicillata*, the black-tufted marmoset, were reported in an urban park in Belo Horizonte, Minas Gerais, Brazil. The epizootic was investigated in collaboration with CETAS/IBAMA and the Zoonoses Department of Belo Horizonte. Nine marmoset carcasses, and 4 sick marmosets were found in the park; the latter exhibited severe neurological symptoms and systemic illness before succumbing within 48 hours. Carcasses were tested for rabies virus and were all negative, and necropsy findings revealed widespread organ damage. In addition, the samples were tested for yellow fever virus, with negative results. Finally, molecular testing, viral isolation and phylogenetic analysis demonstrated human herpesvirus 1 (HHV-1) as the causative agent. The likely source of infection was human-to-marmoset transmission, facilitated by close interactions such as feeding and handling. This study highlights the risks of pathogen spillover between humans and nonhuman primates, emphasizing the need for enhanced surveillance and public awareness to mitigate future epizootics.

## 1. Introduction

Human activities in sylvatic environments and contact with animals, such as non-human primates (NHP), increase the risk of bidirectional pathogen transmission between NHPs and humans. There is a long history of infectious diseases spreading from NHPs to humans, but transmission in the opposite direction has also been observed, including Zika virus, SARS-CoV-2, measles, influenza virus, hepatitis virus, herpesviruses, and others (1– 6). The breakdown of natural barriers between NHPs and humans poses health risks for a much larger population, including humans living in urban areas (4) and NHPs living in pristine forests. In this context, urban parks serve as key points of interaction between humans and free-living NHPs, facilitating cross-infections (6). Urban parks are important components of modern human society, enabling outdoor recreation and alleviating urban heat island effects (7). In many urban parks in the neotropics, marmosets perform essential ecological functions, such as seed dispersal and insect predation, thereby contributing to the regeneration of native plants and to pest control (8). Additionally, their presence can enhance local biodiversity, promoting ecological balance (8,9).

Between 2016 to 2018 Brazil faced significant yellow fever (YF) outbreaks, with over 2,000 human and NHP epizootic cases each, mainly in Minas Gerais state (MG) (10–15). Belo Horizonte (BH), the capital of the state, was also affected, with yellow fever virus (YFV, *Orthoflavivirus flavi*) - positive marmosets’ carcasses being found in different areas of the city, including urban parks (14). Since the outbreaks, cases of monkeys and humans infected with YFV have been continuously reported outside the Amazon Basin in Brazil, including MG state (10,16) and the metropolitan region of BH (17). More recently between March to June 2024, epizootics took place in an urban park in BH, causing the deaths of several black-tufted marmosets and drastically reducing their population in the affected region. Herein, we report the outcomes of our investigation to identify the pathogen responsible for the outbreak.

## 2. Materials and Methods

### 2.1 Study Area and Samples

Belo Horizonte (BH) is in a region of transition between the Atlantic Forest and the Cerrado (Brazilian savannah) habitats (18) in MG, southeast Brazil. Atlantic Forest and Cerrado biomes are among the most biodiverse in the world. Both biomes face threats to their endemic mammal species, many at risk of extinction, making them critical hotspots for conservation (19,20). BH has approximately 80 urban parks (21), which play a crucial role in preserving biodiversity in the face of urban growth. Black-tufted marmosets (*Callithrix penicillata*) live in these parks and other regions of the city, as they are well adapted to urbanized environments. Their ability to exploit diverse habitats, access a variety of food sources, and maintain a versatile diet contributes to their successful adaptation (22–25). The urban park Primeiro de Maio is in the northern part of BH, covering an area of 3.3 hectares (26). It features nine springs, various species of trees, and vertebrate animals including turtles, fish, small mammals, several bird species, and marmosets. The park has different recreational areas and is surrounded by residential areas along its entire perimeter. Park rangers reported that marmosets are attracted to humans, and it is common to observe visitors or neighbors feeding the marmosets.

In March 2024, the first observations of dead and sick marmosets in Primeiro de Maio park were reported, and the epizootic continued until May 2024. A total of 13 affected animals were observed, nine carcasses of marmosets were found at the park and four sick marmosets were also found. In June of 2024, fieldwork and baiting activities were conducted over three days in the park, followed by one day dedicated to trapping and sample collection. During this time only two marmosets were sighted, with one of them being captured and sampled. Park rangers reported that most marmosets in the park were affected by the epizootic. The outbreak was investigated jointly with the Department and the Laboratory of Zoonoses of BH (LZOON-BH), and the Wildlife Rescue Center (CETAS-BH). Four marmosets (two adult males, one infant male and one adult female), found sick in the park Primeiro de Maio were forward to the Wildlife Center of BH (CETAS-BH). They presented fever, apathy, weakness, dehydration, and neurological involvement characterized by multiple seizures, unilateral ptosis, pupil contraction, incoordination, disorientation, and lateral recumbency. Only one individual exhibited ulceration on the lingual mucosa, and all of them died within 48 hours. Following their death, the carcasses of four marmosets (NHP 1342 to 1345), were sent to LZOON-BH and Virus Lab-UFMG for further investigation. At LZOON-BH all carcasses (n=13) tested negative for rabies, by routine protocols. At Virus Lab-UFMG, necropsy was performed, and liver, kidney, heart, lung, tongue, bladder, spleen, testicle and blood clot samples were collected and preserved at −80°C and used for the following analyses.

### 2.2 Virus investigation

Tissue fragments from marmoset carcasses were separately used for total nucleic acid extraction. We performed total viral DNA and RNA extraction using the QIAamp cador Pathogen Mini Kit protocol (QIAGEN, Venlo, Netherlands) on liver, blood clot, lung, kidney, testicle, and spleen samples (approximately 30 mg). These samples were then investigated for YFV and human herpesviruses. Further samples (serum, heart, intestine and bladder, tested only for herpesviruses) had total DNA extracted with High Pure Viral Nucleic Acid Kit (Roche, Basel, Switzerland). Mechanical tissue disruption was performed using three 2 mm borosilicate beads and a bead beater (Mini-Beadbeater-16, BioSpec Products, Bartlesville, OK, USA). Samples were extracted in batches of up to 14 samples plus a negative extraction control (nuclease-free water).

RT-qPCR was performed using the GoTaq® Probe 1-Step RT-qPCR System (Promega) to assess RNA viability through the endogenous control β-actin gene (Table S1) (27), confirming all samples as suitable for analysis. Total viral RNA/DNA obtained from liver, blood clot, lung, kidney, testicle, and spleen (2.5 µL) along with negative extraction controls were screened in duplicate for YFV RNA, targeting the 5’ untranslated region of the viral genome (Table S1) (28). In each set of RT-qPCR a non-template control (nuclease-free water) and a positive control (YFV 17DD RNA) was included. Samples were also tested for human herpesviruses 1 and 3 (HHV-1/2, and HHV-3) by qPCR (GoTaq® qPCR Promega Corporation, Madison, WI, USA), using primers targeting the polymerase gene (29) Table S1). To distinguish HHV-1 from HHV-2, samples were tested using primers targeting the glycoprotein D (gD) gene of HHV-1 and glycoprotein G (gG) gene of HHV-2 (30) (Table S1). Extracted DNA from HHV-1 and HHV-3 were used as positive control, as well as a non-template control (nuclease-free water) in each qPCR and PCR reaction.

A fragment of the tongue with a lesion from NHP 1342 was macerated (50 mg of tissue in 200 µL of Eagle’s minimal essential medium (MEM) without fetal bovine serum (FBS) by mechanical disruption as described above. After maceration, the product was centrifuged for 10 minutes at 10,000 x g and used as an inoculum on the Vero cells monolayer. Vero cells cultivated in 12-well plate (1.8 × 10_5_ Vero cells per well) were inoculated with 180 µl of the 1:2 diluted macerated and maintained in MEM supplemented with 5% FBS at 37°C in a 5% CO_2_ incubator. Cells were observed in inverted microscope and after three days of infection.

The supernatant of infected cells was collected and subjected to DNA extraction and PCR targeting gD (HHV-1) and gG (HHV-2) genes (30) and conventional PCR targeting the glycoprotein B (gB) gene, generating a 400 nt amplicon (31) (Table S1). PCR amplicon was analyzed by electrophoresis in an 8% polyacrylamide gel, stained with SYBR Gold (Invitrogen – Thermo Fisher Scientific, EUA) and examined under a Blue-Light Transilluminator (Safe Imager™ 2.0 Blue-Light Transilluminator, Invitrogen – Thermo Fisher Scientific, EUA). The observed amplicon was purified and sequenced by dideoxy method with the ABI3130 platform (Applied Biosystems, USA). The consensus sequence was generated (Chromas software version 2.6.6) and the Basic Local Alignment Search Tool (BLAST) was used to find regions of local similarity between sequences on nucleotide databases. Consensus sequence was aligned with 14 sequences (GenBank accession numbers in Supplementary file) in MEGA12 (https://www.megasoftware.net/) using Tamura-Nei (1993) model and the Maximum-likelihood method with 1000 bootstrap replicates.

## 3. Results and Discussion

Considering the persistent circulation of YFV in MG, including the metropolitan region of BH (10, 17, 32-34), we initially investigated YFV infection in carcasses. Samples were negative, indicating no active infection with YFV. Brains were removed in LZOON-BH for rabies investigation and were found to be negative for rabies virus.

The four necropsied carcasses of marmosets were in an advanced stage of autolysis, but it was possible to note extensive areas of multifocal hepatic necrosis, characterized by pale-colored zones with hemorrhagic centers, indicative of cellular degeneration (35-36). The lungs showed foci of interstitial pneumonia with consolidation, diffuse edema, and hemorrhage, consistent with the severe dyspnea. Brains were removed in LZOON-BH for rabies investigation and were not available for analysis. The regional lymph nodes were enlarged and hyperemic, suggesting an intense systemic inflammatory response (Table S2). The tongue of NHP 1342 exhibited well-defined necrotic ulcers with a hemorrhagic halo, associated with intense sialorrhea and dysphagia (Figure 1A).

**Figure 1.**
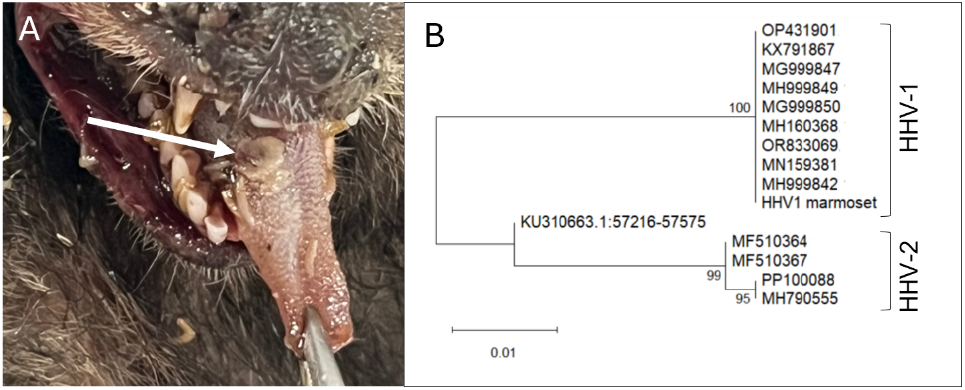
Human herpesvirus 1 in marmosets, during an epizootic in Belo Horizonte, Minas Gerais, Brazil. (A) Marmoset tongue presenting a herpes lesion. (B) Evolutionary analysis inferred using the Maximum Likelihood method and Tamura-Nei (1993) model. The percentage of replicate trees in which the associated taxa clustered together (1.000 replicates) is shown next to the branches. The analyses included 15 partial sequences of glycoprotein B coding gene with 351 positions in the final dataset. Evolutionary analyses were conducted in MEGA12.

Next, we investigated the marmosets for human herpesvirus infection. Liver, lung, kidney, testicle, heart, intestine, or bladder samples were simultaneously tested for HHV-1/2 and HHV-3. While samples from the 4 marmosets were negative for HHV-3, they were positive for HHV-1/2 by qPCR (Table 1, Table S3). Viral isolation was conducted using a fragment of the lesion on the tongue (marmoset 1342). After three days of infection, cell rounding, detachment, and lysis across most of the cell monolayer was observed in infected cells compared to the control cells. The supernatant was collected and subjected to DNA extraction, diluted 1:2, and tested positive for HHV-1 (Cq = 12,5), and phylogenetic analysis using partial sequence of gB gene (Genbank accession number PV358090) confirmed the infection with HHV-1 (Figure 1B). Serum from the free-living marmoset captured in June/2024 (CT-24-203) in the park was tested by qPCR and negative for HHV-1 (Table 1).

**Table 1.**
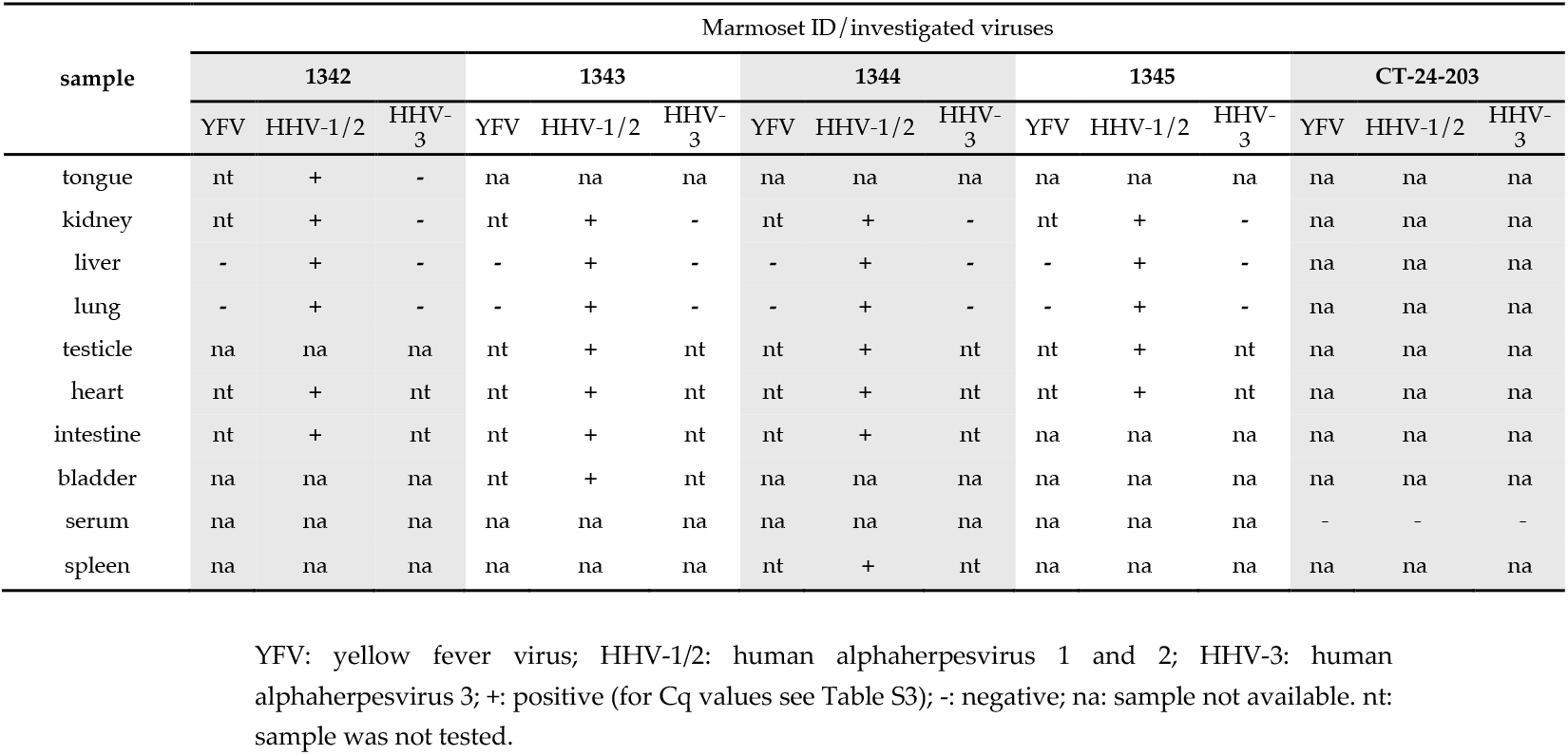
Molecular investigation of yellow fever virus and human herpesviruses in *Callithrix penicilatta*.

| sample | Marmoset ID / investigated viruses |  |  |  |  |  |  |  |  |  |  |  |  |  |  |
| --- | --- | --- | --- | --- | --- | --- | --- | --- | --- | --- | --- | --- | --- | --- | --- |
|  | 1342 |  |  | 1343 |  |  | 1344 |  |  | 1345 |  |  | CT-24-203 |  |  |
|  | YFV | HHV-1/2 | HHV-3 | YFV | HHV-1/2 | HHV-3 | YFV | HHV-1/2 | HHV-3 | YFV | HHV-1/2 | HHV-3 | YFV | HHV-1/2 | HHV-3 |
| tongue | nt | + | - | na | na | na | na | na | na | na | na | na | na | na | na |
| kidney | nt | + | - | nt | + | - | nt | + | - | nt | + | - | na | na | na |
| liver | - | + | - | - | + | - | - | + | - | - | + | - | na | na | na |
| lung | - | + | - | - | + | - | - | + | - | - | + | - | na | na | na |
| testicle | na | na | na | nt | + | nt | nt | + | nt | nt | + | nt | na | na | na |
| heart | nt | + | nt | nt | + | nt | nt | + | nt | nt | + | nt | na | na | na |
| intestine | nt | + | nt | nt | + | nt | nt | + | nt | na | na | na | na | na | na |
| bladder | na | na | na | nt | + | nt | na | na | na | na | na | na | na | na | na |
| serum | na | na | na | na | na | na | na | na | na | na | na | na | - | - | - |
| spleen | na | na | na | na | na | na | nt | + | nt | na | na | na | na | na | na |

YFV: yellow fever virus; HHV-1/2: human alphaherpesvirus 1 and 2; HHV-3: human alphaherpesvirus 3; +: positive (for Cq values see Table S3); -: negative; na: sample not available. nt: sample was not tested.

Both NHPs and humans host a variety of specific herpesviruses. While these viruses typically cause asymptomatic infections in their natural hosts, they can lead to severe disease when transmitted to different species (35). For example, herpesvirus B (*Simplexvirus macacinealpha1*) is rarely responsible for disease in its natural host, the macaque, but infection of humans results in a disseminated viral infection characterized by ascending paralysis and a high case fatality rate. The opposite situation can also occur when HHV-1/2 and HHV-3 infects NHPs (35,36). Although HHV-1 infection in old world monkeys usually results in mild, self-limiting oral vesicular lesions, New World monkeys usually develop lethal disseminated disease. In NHPs, HHV-1 is commonly responsible for oral lesions and encephalitis in adults, whereas HHV-2 is usually a sexually transmitted disease causing a genital infection in adults and a disseminated infection in infants (36,37).

The observed lesion pattern and clinical signs, (1-4, 36) and molecular testing confirmed systemic infection by HHV-1. NHPs are not naturally infected by HHV-1 in the wild; thus, infection occurs through contact with infected humans or contaminated fomites (2,3, 36). Once established in a population of New World NHPs, HHV-1 spreads rapidly, potentially leading to high mortality and morbidity within the population (3), as observed in the epizootic studied here. Unfortunately, we were not able to track the exact source of HHV-1 infection in the marmosets in our study. However, given that close contact between NHP and humans is a key factor in pathogen transmission (3,24,38,39) and frequent interactions - such as physical contact and feeding-between park’s visitors and neighbors - are reported by park rangers, probably humans were the source of HHV-1 infection in NHPs.

Direct and indirect interactions between humans and NHP increase the risk of interspecific disease transmission, posing challenges for both species conservation and public health (40-41). Beyond these risks, excessive proximity to black-tufted marmosets can lead to significant behavioral changes, making them dependent on humans for food, which has negative consequences for their health (24,42) This dependency can weaken their ecological roles in pest control and seed dispersal, reduce their natural antipredator responses, and lower their aversion to human presence (43-45). As a result, human-marmoset interactions may escalate into conflicts and, in extreme cases, acts of aggression toward people (38,24).

Monitoring NHP infections is crucial for understanding transmission dynamics, identifying pathogen persistence, and detecting potential spillover or spillback events within disease cycles. Addressing this complex issue requires an integrated approach that combines surveillance efforts with targeted educational initiatives. Promoting responsible human behavior toward wildlife is essential for mitigating the risks of viral spillback and preventing future epizootics.

## Supporting information

Supplementary Tabels

## Supplementary Materials

The following supporting information can be downloaded at: www.mdpi.com/xxx/s1, Table S1: Primers and probes used for molecular investigation of yellow fever virus and human herpesviruses; Table S2: Necropsies findings in *Callithrix penicilatta* carcasses; Table S3: Results of yellow fever virus (RTqPCR) and Human Herpesviruses (qPCR) molecular investigation in *Callithrix penicilatta*.

## Author Contributions

Conceptualization, B.P.D.; Methodology, B.P.D., G.F.G.-O., M.F.B.; Formal Analysis, G.F.G.-O., M.F.B, and B.P.D.; Investigation, G.F.G.-O., M.F.B., D.J.d.C., M.A.O., A.M.O.P.; Resources, M.F.B., D.J.d.C., M.A.d.O., A.M.d.O.P., P.A.A., C.B., D.A.d.R.V., É.P.T.T., T.L.S., T.M.M., M.C.S., M.G.A., I.V.S., É.M.d.M., F.E.N.d.G.; Writing—Original Draft Preparation, G.F.G.-O., M.F.B, A.M.O.P., and B.P.D.; Writing—Review and Editing, G.F.G-O., M.F.B., D.J.d.C., M.A.d.O., A.M.d.O.P., P.A.A., C.B., D.A.d.R.V., É.P.T.T., T.L.S., T.M.M., M.C.S., M.G.A., I.V.S., É.M.d.M., F.E.N.d.G., K.A.H., N.V., B.P.D.; Visualization, G.F.G.-O., and B.P.D.; Supervision, B.P.D.; Project Administration, K.A.H., N.V., and B.P.D.; Funding Acquisition, K.A.H., N.V., and B.P.D. All authors have read and agreed to the published version of the manuscript.

## Institutional Review Board Statement

Ethical approval for the study was obtained from the Committee on Ethics in the Use of Animals at the Universidade Federal de Minas Gerais (protocol CEUA 290/2022), and research in federal conservation units was authorized by the Chico Mendes Institute for Biodiversity Conservation (protocol SISBIO 77400). The project is registered at SISGEN-Brazil under protocol A8091C3.

## Informed Consent Statement

Not applicable.

## Data Availability Statement

All research data is shared within the manuscript and Supplementary Tables 1, 2 and 3 and in ZENODO repository under https://doi.org/10.5281/zenodo.15061612

## Acknowledgments

We would like to thank Secretaria Municipal de Saúde de Belo Horizonte, Laboratório de Zoonoses, and Centro de Controle de Zoonoses da Prefeitura de Belo Horizonte, Department of Zoonoses of Belo Horizonte, Secretaria de Estado de Saúde de Minas Gerais, Centro de Triagem de Animais Silvestres of Belo Horizonte, colleagues from Laboratório de Vírus/UFMG, Pró-Reitorias de Graduação, Pós-graduação, and Pesquisa from Universidade Federal de Minas Gerais/Brazil.

## Abbreviations

The following abbreviations are used in this manuscript:

NHP: Non-Human Primate
IBAMA: Instituto Brasileiro do Meio Ambiente e dos Recursos Naturais Renováveis
CETAS: Centro de Triagem de Animais Silvestres
BH: Linear dichroism
MG: Minas Gerais
YFV: Yellow Fever Virus
HHV: Human Herpes Virus
LZOON: Laboratório de Zoonoses
UFMG: Universidade Federal de Minas Gerais
DNA: Deoxyribonucleic Acid
RNA: Ribonucleic Acid
PCR: Polymerase Chain Reaction
RT-qPCR: Reverse Transcriptase Real Time Polymerase Chain Reaction gD Glycoprotein D
gG: Glycoprotein G
gB: Glycoprotein B
MEM: Eagle’s minimal essential medium
FBS: Fetal Bovine Srum
nt: Nucleotide

