## Supplementary Tabels for "Human herpesvirus 1 associated with epizootics in Belo Horizonte, Minas Gerais, Brazil"

**Garcia-Oliveira et al.**

Table S1: Primers and probes used for molecular investigation of yellow fever virus and human herpesviruses

| Target | Primer or probe | Sequence | Region | Reference |
|---|---|---|---|---|
| YFV | YFallF primer | 5'-GCTAATTGAGGTGYATTGGTCTGC-3' | 5' UTR | Domingo et al, 2012 |
| | YFallR primer | 5'-CTGCTAATCGCTCAAMGAACG-3' | | |
| | YFallP probe | 5′-FAM-ATCGAGTTGCTAGGCAATAAACAC-TMR-3′ | | |
| HHV-1/HHV-2 | HHV-1/2 F | 5′-ATCAGGGTAGCCGCGCGTGTGACA-3′ | DNA POL | De Oliveira, 2015 |
| | HHV-1/2 R | 5′-CATACCGGAACGACCACAC-3′ | | |
| HHV-3 | HHV-3 F | 5′-CATCTGCAATTATGGCTCCAA-3′ | DNA POL | De Oliveira, 2015 |
| | HHV-3 R | 5′-GTTTCCATTGCTGAAT-3′ | | |
| Endogenous control | β-actin F | 5'-CCAACCGCGAGAAGATGA-3' | β-actin | Rezende et al, 2019 |
| | β-actin R | 5'-CCAGAGGCGTACAGGGATAG-3' | | |
| HHV | Primer KS30 | 5'-biotinyl-TTCAAGGCCACCATGTACTACAAAGACGT-3' | Glycoprotein B | Orle et al, 1996 |
| | Primer KS31 | 5'-biotinyl-GCCGTAAAACGGGGACATGTACACAAAGT-3' | | |
| HHV-1 | Forward primer (HSV1UP) | 5'-CGGCCGTGTGACACTATCG-3' | Glycoprotein D | Weidmann et al, 2003 |
| | Reverse primer (HSV1DP) | 5'-CTCGTAAAATGGCCCCTCC-3' | | |
| | Probe (HSV1P) | 5'-CCATACCGACCACACCGACGAACC-3' | | |
| HHV-2 | Forward primer (HSV2UP) | 5'-CGCTCTCGTAAATGCTTCCCT-3' | Glycoprotein G | Weidmann et al, 2003 |
| | Reverse primer (HSV2DP) | 5'-TCTACCCACAACAGACCCACG-3' | | |
| | Probe (HSV2P) | 5'-CGCGGAGACATTCGAGTACCAGATCG-3' | | |

YFV: Yellow Fever Virus – *Orthoflavivirus flavi*; HHV: Human Herpesvirus; HHV-1: human alphaherpesvirus 1 - *Simplexvirus humanalpha1*; HHV-2: human alphaherpesvirus 2 - *Simplexvirus humanalpha*2; HHV-3: Varicela-Zoster virus - *Varicellovirus humanalpha3;* 5′UTR: 5' untranslated region; DNA POL: DNA polymerase.

Table S2: Necropsies findings in *Callithrix penicilatta* carcasses

| | Marmosets/Macroscopic aspects of necropysis | | | | | | | | | | | | | | | |
|---|---|---|---|---|---|---|---|---|---|---|---|---|---|---|---|---|
| | **1342** | | | | **1343** | | | | **1344** | | | | **1345** | | | |
| **Sample** | **Coloration** | **Size** | **Consistency** | **Aspect** | **Coloration** | **Size** | **Consistency** | **Aspect** | **Coloration** | **Size** | **Consistency** | **Aspect** | **Coloration** | **Size** | **Consistency** | **Aspect** |
| brain | na | na | na | na | na | na | na | na | na | na | na | na | na | na | na | na |
| heart | reddish | enlarged | hardened | rough | normal | normal | soft | smooth | darkened | enlarged | ni | ni | darkened | normal | hardened | normal |
| lung | normal | normal | soft | smooth | reddish | enlarged | hardened | rough | reddish | enlarged | flabby | hemorrhagic | reddish | normal | flabby | hemorrhagic |
| liver | yellowish | reduced | flabby | necrotic | darkened | reduced | necrotic | flabby | darkened | enlarged | ni | ni | na | na | na | na |
| kidney | darkened | enlarged | hardened | granular | normal | normal | soft | smooth | reddish | normal | soft | smooth | anemic | enlarged | soft | normal |
| spleen | anemic | normal | normal | smooth | greenish | enlarged | flabby | rough | na | na | na | na | na | na | na | na |
| stomach | normal | normal | soft | smooth | yellowish | normal | smooth | soft | na | na | na | na | na | na | na | na |
| intestine | greenish | enlarged | flabby | rough | normal | reduced | rough | hardened | reddish | enlarged | ni | hemorrhagic | na | na | na | na |

na: samples not available; ni: aspect not identified.

Table S3: Results of yellow fever virus (RTqPCR) and Human Herpesviruses (qPCR) molecular investigation in *Callithrix penicilatta*

Marmosets/investigated viruses (and Cq values when samples were positive)

| **sample** | **1342** | | | **1343** | | | **1344** | | | **1345** | | |
|---|---|---|---|---|---|---|---|---|---|---|---|---|
| | YFV | HHV-1/2 | HHV-3 | YFV | HHV-1/2 | HHV-3 | YFV | HHV-1/2 | HHV-3 | YFV | HHV-1/2 | HHV-3 |
| TONGUE | nt | + / 27.6 | - | na | na | na | na | na | na | na | na | na |
| KIDNEY | nt | + / 31 | - | nt | + / 34.3 | - | nt | + / 30.5 | - | nt | + / 33.2 | - |
| LIVER | - | + / 34.4 | - | - | + / 37.8 | - | - | + / 31.9 | - | - | + / 34.3 | - |
| LUNG | - | + / 35.2 | - | - | + / 37.5 | - | - | + / 33.8 | - | - | + / 29.9 | - |
| TESTICLE | na | na | na | nt | + / 35.2 | nt | nt | + / 32 | nt | nt | + / 29.3 | nt |
| HEART | nt | + / 26.5 | nt | nt | + / 34 | nt | nt | + / 30 | nt | nt | + / 26 | nt |
| INTESTINE | nt | + / 32.5 | nt | nt | + / 31.5 | nt | nt | + / 29 | nt | na | na | na |
| BLADDER | na | na | na | nt | + / 31 | nt | na | na | na | na | na | na |
| SPLEEN | na | na | na | na | na | na | nt | + / 34.7 | nt | na | na | na |

YFV: yellow fever virus; HHV-1/2: human alphaherpesvirus 1 and 2; HHV-3: Varicella zoster virus/human alphaherpesvirus 3; +: positive (for Cq values see Table S3); -: negative; na: sample not available. nt: sample was not tested. Cq:cycle threshold in real-time PCR
